# Bridging theories for ecosystem stability through structural sensitivity analysis of ecological models in equilibrium

**DOI:** 10.1101/2019.12.24.887901

**Authors:** Jan J. Kuiper, Bob W. Kooi, Garry D. Peterson, Wolf M. Mooij

## Abstract

Ecologists are challenged by the need to bridge and synthesize different approaches and theories to obtain a coherent understanding of ecosystems in a changing world. Both food web theory and regime shift theory shine light on mechanisms that confer stability to ecosystems, but from a different angle. Empirical food web models are developed to analyze how equilibria in real multi-trophic ecosystems are shaped by species interactions, and often include linear functional response terms for simple estimation of interaction strengths from observations. Models of regime shifts focus on qualitative changes of equilibrium points in a slowly changing environment, and typically include non-linear functional response terms. Currently, it is unclear how the stability of an empirical food web model, expressed as the rate of system recovery after a small perturbation, relates to the vulnerability of the ecosystem to collapse. Here, we conduct structural sensitive analyses of classical consumer-resource models in equilibrium along an environmental gradient. Specifically, we change non-proportional interaction terms into linear ones, while maintaining the equilibrium biomass densities and flux of matter, to analyze how alternative model formulations shape the stability properties of the equilibria. The results reveal no consistent relationship between the stability of the original models and the linearized versions, even though they describe the same biomass values and material flows. We use these findings to discuss whether stability analysis of observed equilibria by empirical food web models can provide insight into regime shift dynamics, and highlight the challenge of bridging alternative modelling approaches in ecology and beyond.

## INTRODUCTION

Human action is transforming global ecosystems through processes such as land conversion, climate change, overexploitation and pollution (Millenium Ecosystem Assessment 2005). These changes in structure and function alter the contributions of nature to people (IPBES 2018). Changes in ecosystem functioning are particularly troublesome for people when they are large, abrupt and difficult to reverse (Scheffer et al. 2001). Such catastrophic transitions in ecosystem state have been observed in various ecosystems including shallow lakes, peat bogs, coral reefs and arid range lands (Rocha et al. 2015). A principal challenge for ecologists is to understand and forecast the stability properties of natural ecosystems in a rapidly changing world (Clark et al. 2001; Evans 2012).

Assessing the stability properties of complex ecosystems requires more than a single metric (Peterson et al. 1997; Ives and Carpenter 2007; Donohue et al. 2013). One classical measure is the rate at which a stable system recovers from a small perturbation, originally defined as resilience by Pimm (1984) and later referred to as engineering resilience by Holling (1996). Importantly, dynamical systems theory predicts that this engineering resilience may be used as an indicator of the magnitude of disturbance that a system can absorb without undergoing radical change in its structure, that is, ecological resilience (Holling 1996; Van Nes and Scheffer 2007; Boettiger et al. 2013). This principle underlies the development of statistical techniques that may detect signals of ‘critical slowing down’ in high frequency time series, potentially providing early warning to abrupt regime shifts (Scheffer et al. 2009). Unfortunately, this statistical approach is still in its infancy (Spears et al. 2017), and many limitations remain (Dakos et al. 2015). Alternatively, dynamical models may be used to understand and predict the resilience of ecological systems. A conventional method is local stability analysis of the Jacobian matrix of all first-order partial derivatives. The real part of the dominant eigenvalue of the Jacobian matrix evaluated at the equilibrium reveals the rate at which a small perturbation to a system in equilibrium decays (Pimm and Lawton 1977; Neubert and Caswell 1997). This recovery rate can be considered a measure of engineering resilience (Van Nes and Scheffer 2007). Ideally it would be possible to construct a mathematical description of a natural ecosystem and evaluate its local stability using the Jacobian matrix to obtain a reliable estimate of the actual engineering resilience of the system that is portrayed, ultimately to be used as an indicator of ecological resilience by ecosystem managers.

The modelling of species interactions often forms the first step in analyzing more complex behavior of ecosystems. As such, the seminal work by Lotka (1920) and Volterra (1926), who independently of each other discovered the cycles that arise in a set of coupled differential equations representing consumers and resources, forms the foundation for innumerable ecological models that have been developed since. The Lotka-Volterra (LV) system is one of the earliest models in mathematical ecology and represents the simplest model of predator-prey interactions, using proportional (linear) per capita growth and mortality rates and mass-action interaction rates. To describe and analyze more realistic ecological systems the classical LV system can be extended along two fundamentally different complexity axes: (i) the number of interacting species and their arrangement in a network, and (ii) the type of functional response terms to characterize the interactions.

Minimal dynamic models, which consist of just a few equations, are the pre-eminent tools for studying the often striking and surprising effects of incorporating non-proportional interaction terms on model outcome (Wangersky 1978; Kooi 2003). One of the most famous examples comes from Rosenzweig and MacArthur (1963), who replaced the linear functional response interaction term of the original LV model with a Holling type II functional response, and included logistically instead of proportionally growing resource. Their observation that increasing resource density tends to destabilize the system led to the formulation of the ‘paradox of enrichment’ (Rosenzweig 1971). Minimal models are often used to scrutinize the local stability along a gradient of environmental change to detect bifurcations that reveal qualitative changes in the long-term dynamics, e.g. productivity in the Rosenzweig-MacArthur example (Kooi 2003). As such, minimal dynamical models help to unveil which ingredients are minimally required to evoke phenomena that are qualitatively similar to phenomena observed in real life (Scheffer 2004). For example, minimal models have been decisive in revealing positive feedback loops as a key ingredient for the emergence of alternative stable states and the occurrence critical regime shifts (May 1977; DeAngelis et al. 1986; Scheffer 1989). These simple models thus provide a strong theoretical foundation for the concept of ecological resilience (Holling 1973; Scheffer et al. 2001; Van Nes and Scheffer 2007), which presumes the existence of multiple alternate states in nature. However, minimal models of ecosystems are not intended to give quantitative descriptions of the system under study. Hence, minimal models are not suitable for producing accurate estimations of the stability properties of observed complex ecosystems.

Potentially more suitable candidates are empirical food web models with numerous coupled equations, which provide some of the most detailed mathematical descriptions of natural ecosystems to date. We here focus on a particular class of food web models that have been developed specifically for local stability analysis of observed equilibria in multi-trophic ecosystems (De Ruiter et al. 1995; Moore and de Ruiter 2012). Interestingly, despite the important theoretical results on stability that were obtained by entering non-linear terms, these empirically-based food web models tend to adhere to the use of Lotka-Volterra type equations with linear interspecific interaction terms (Neutel and Thorne 2016). The crux is that linear interaction terms make it possible to derive the partial derivatives of Lotka-Volterra type growth equations in equilibrium directly from readily available empirical information such as biomass densities and feeding rates (De Ruiter et al. 1993, 1995; Neutel and Thorne 2016; Heijboer et al. 2017). The partial derivatives represent per capita interaction strengths and give the elements of the Jacobian matrix representation of the food web. Typically the dominant eigenvalue of the Jacobian matrix is used as a dependent variable for comparing systems with different network structures to identify which network properties are relatively important for stability (Van Altena et al. 2014). Studies unravelling the ‘stability in real food webs’ have yielded compelling insights into which stabilizing structures are prevalent in nature and hence should be preserved (De Ruiter et al. 1995; Neutel et al. 2002; Jacquet et al. 2016).

Unfortunately, however, these studies have not clearly explained how the calculated stability in multi-trophic ecosystems relates to the concept of ecological resilience and the imminence of catastrophic regime shifts. Local (fixed-point) stability analysis of empirically-based food web models emphasizes the bouncing back after disturbance, and thus the engineering view on resilience (Pimm 1984). An advantage is easy comparison with empirical ecological research in, for example, community ecology, where recovery time after a natural or experimental disturbance might be used to detect relationships between biodiversity and stability (Tilman and Downing 1994; McCann 2000; Loreau et al. 2001; Kuiper et al. 2014). Insights may also be translated to other disciplines that focus on disturbance and recovery, such disaster risk management and economic geography (Fingleton et al. 2012), which often have an implicit focus on resisting and controlling change. However, although seldom articulated, there is also an implicit assumption of global stability (Gunderson 2000), that is, that there is only a single stable interior state (all species with positive biomasses) showing a attracting equilibrium, limit cycle or chaos. Indeed, the few studies that have analyzed empirical food web models along environmental gradients of e.g. productivity (Neutel et al. 2007), grazing pressure (Andres et al. 2016) and climate warming (Schwarz et al. 2017) did not discuss their results in relation to ecological resilience and potential whole-system regime shifts. Kuiper et al. (2015), however, did find evidence for a relationship when they parameterized food web models using data sampled from a virtual reality created by a complex ecosystem model with alternative stable states, but the complexity of their models makes it difficult to fully grasp the results. As a result, **it remains largely unclear to what extent the stabilizing structures in trophic networks that are identified through local stability analysis of empirical food web models are important for preventing catastrophic regime shifts (Kuiper et al. 2015)**. Reconciling food web theory and regime shift theory is crucial for better understanding the relation between biodiversity decline and ecosystem collapse (Cardinale et al. 2012; Downing et al. 2012), and the development of empirical indicators of ecological resilience to anticipate regime shift.

To serve as an indicator of ecological resilience and reveal insights in the vulnerability of the real-world ecosystem to regime shifts, the calculated stability of empirically-based food web models must correspond reasonably well to the actual engineering resilience of the system that is being modelled (Figure 1; Van Nes and Scheffer 2007; Aldebert et al. 2016a; Neutel and Thorne 2016). A fundamental underlying question is how the assumption of linear interaction terms shapes the calculated stability properties of empirically-based food web models, and potentially disrupts the connection with the stability properties of the actual ecosystem that is being modelled, considering that it is common knowledge that the type of functional response term can have a drastic influence on the stability properties of a dynamic model.

**Figure 1.**
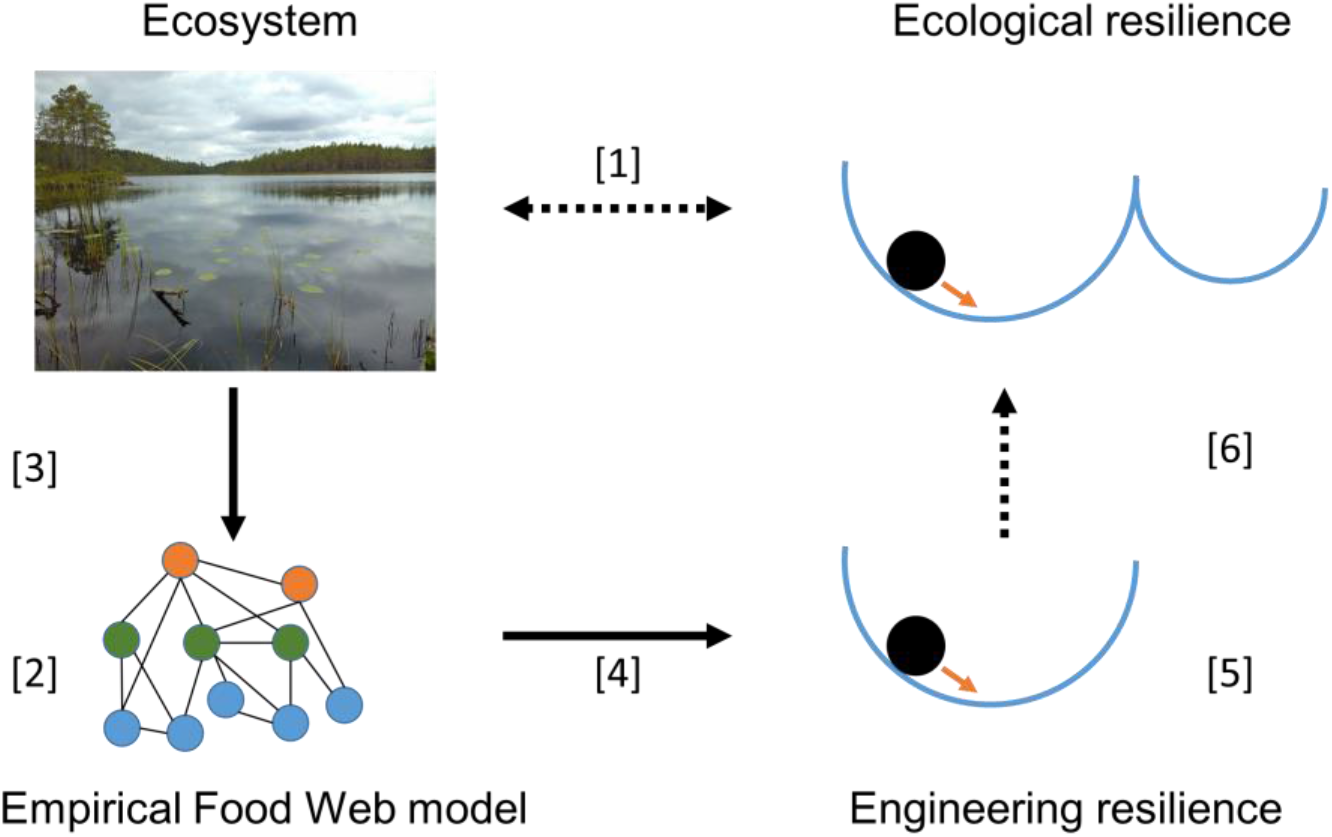
A challenge for ecologists is to provide human societies with estimates of how much stress an ecosystem can withstand without losing the ability to recover and change to an alternate state, that is, ecological resilience [1]. Empirically-based food web models are developed to analyze the stability of complex ecosystems [2]. Empirical observations are used to construct a detailed material flux description of the ecosystem [3]. When equilibrium conditions and type I functional response terms are assumed, this food-web description can be used to estimate interaction strengths and quantify the elements of the Jacobian matrix representation of the food web model. Local (fixed-point) stability analysis of the Jacobian matrix provides the stability properties of the model system [4]. The dominant eigenvalue provides an estimate of the return rate equilibrium following a perturbation and hence can be used as a measure of engineering resilience [5]. It is unclear whether this measure of engineering resilience provides a meaningful indicator of ecological resilience [6] of the real-world ecosystem that is being portrayed [1]. Photo credit: the corresponding author.

In this paper we address this question by conducting structural sensitive analyses of classical consumer-resource models in equilibrium along an environmental gradient. Specifically, we change non-proportional interaction terms into linear ones, while maintaining the equilibrium biomass densities and flux of matter, to analyze how alternative model formulations shape the stability properties of the equilibria. We repeat our analysis along an environmental gradient, knowing that noticeable qualitative changes in model behavior will occur in the more complex systems with non-linear interaction terms. We check to what extent the stability properties of these alterative model systems with identical equilibrium solutions correspond, and evaluate whether the stability of the simpler systems with linear interaction terms may serve as an indicator of the ecological resilience of the more complex non-linear systems.

While the historical trend has been to add complexity to Lotka-Volterra models, we argue that there is much to gain from starting with complex models and working from them towards simpler models. This may help to identify the least amount of information needed to disclose the stability properties of complex ecosystems, and better understand what mathematical models can teach us about the stability properties or real systems. Yet, we deliberately use minimal dynamic models to benefit from their analytical tractability and explain our findings mathematically. We then discuss our results in light of empirical food web models with linear interaction terms that have been used for stability analysis of observed equilibria in real ecosystems that are inherently complex and non-linear, and identify research needs to reconcile food web theory and regime shift theory.

## METHODS

We base our theoretical experiments on three widely used minimal models of consumer-resource interactions with non-linear functional response terms. For each of the models we present explicit expressions for the equilibria and the elements of the Jacobian matrix. The elements of the matrix determine the dynamic behaviour of the system in the vicinity of the equilibrium and are used to calculate the eigenvalues. Subsequently, we produce linear versions of the original models, by changing the non-proportional functional response terms into proportional terms, while maintaining the equilibrium solutions. We then parameterize the original non-linear models to obtain values of the equilibrium solutions and use these equilibrium values to parametrize the linear counterpart models. This enables us to analyse the local stability of the equilibria using both the non-linear and the linear model systems and compare the stability properties. We repeat this step along a gradient of environmental stress, knowing that bifurcations, characterized by qualitative changes in model behaviour, will occur in the non-linear systems. We check whether the stability analysis of the simpler system discloses any information about the changing stability properties occurring in the complex non-linear system.

### The models

We focus on three key extensions of the classical Lotka-Volterra (LV) consumer-resource model. We define all models for algae and zooplankton, and, therefore, identify the resource as *A* and the consumer as *Z* (Table 1). The original LV model contains proportional growth and death rates of resource and consumer, respectively, and a mass-action interaction term between consumer and resource, that is, a Holling type I functional response without ceiling. The dynamics of the original LV model are given by:

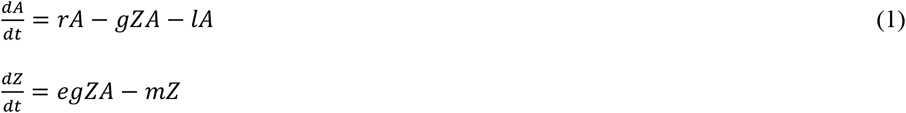

**Table 1.** States and parameters of the consumer-resource models and their linearized counterpart models.

|  | Description | Unit | Value |
| --- | --- | --- | --- |
| $A$ | Resource | Resource unit | |
| $Z$ | Consumer | Consumer unit | |
| $r$ | Maximum growth rate resource | Time<sup>-1</sup> | 0.5 |
| $K$ | Carrying capacity resource | Resource unit | 0-10 |
| $g$ | Maximum intake rate consumer | LV and LVV model:<br>consumer unit<sup>-1</sup>·time<sup>-1</sup>;<br>RM and RMS model:<br>resource unit·consumer unit<sup>-1</sup>·time<sup>-1</sup> | 0.4 |
| $a$ | Half saturation consumer on resource | Resource unit | 0.6 |
| $l$ | Resource loss rate | Time<sup>-1</sup> | 0.01 |
| $e$ | Consumer-resource conversion efficiency | Consumer unit·resource unit<sup>-1</sup> | 0.6 |
| $F$ | Maximum intake rate top-consumer | Consumer unit·time<sup>-1</sup> | 0-0.3 |
| $z$ | Half saturation rate top-consumer on consumer | Consumer unit | 0.5 |
| $m$ | Consumer loss rate | Time<sup>-1</sup> | 0.15 |
| $r_{lin}$ | Growth rate resource | Time<sup>-1</sup> | |
| $g_{lin}$ | Intake rate consumer | Time<sup>-1</sup> | |
| $F_{lin}$ | Intake rate top-consumer | Time<sup>-1</sup> | |

The expressions of the equilibria are:

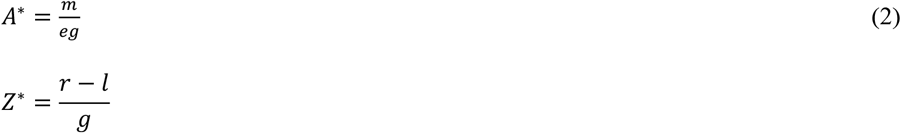

The expressions of the elements of the Jacobian matrix are:

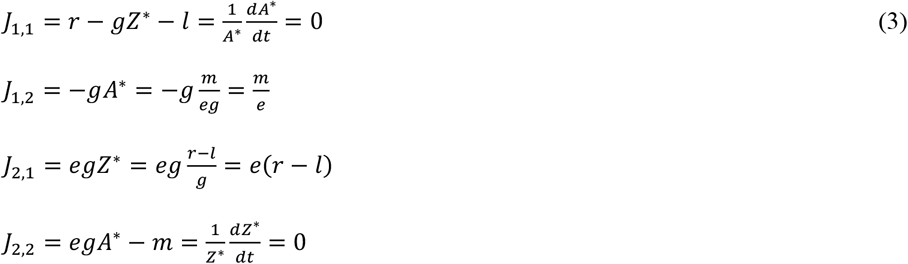

Following ecological modelling terminology, in this paper we will refer to consumer-resource models with proportional growth, loss and interaction terms as having ‘linear’ equations, and refer to models that have non-proportional terms as ‘non-linear’ equations (cf. Arditi and Michalski 1996).

#### The Lotka-Volterra-Verhulst model

The first key extension of the original LV model is the inclusion of negative density dependence in the resource so as to prevent unbounded growth in absence of the consumer. A classical formulation of negative density dependence is the logistic equation introduced by Verhulst (Verhulst 1845). Hence, we replace the linear growth term *rA* in the resource with a the non-linear logistic term 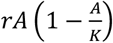, and refer to this extension as the Lotka-Volterra-Verhulst (LVV) model. The dynamics of the LVV model are given by:

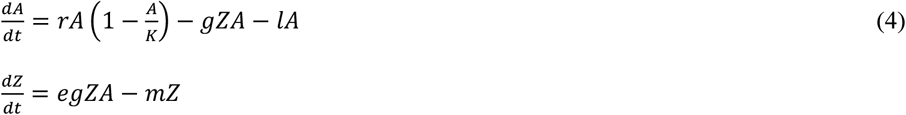

The expressions of the equilibria are (for Z^∗^ > 0):

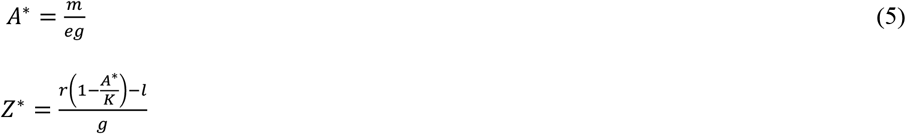

The only element of the Jacobian Matrix that is affected by the addition of the logistic growth term is *J*_*1,1*_. The partial derivative of the logistic term is 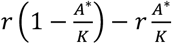, which is included in *J*_*1,1*_ (compare Eq. 3). The full expression of the element is:

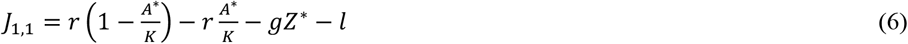

#### The Rosenzweig-MacArthur model

A second key extension of the LV model results from the realization that consumers approach a maximum intake rate at high resource levels. This can for instance be achieved by replacing the linear interaction term of the Lotka Volterra model gZA with a non-linear Holling type II functional response 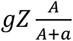 (Holling 1959), which in turn is mathematically equal to Michaelis-Menten kinetics (Michaelis and Menten 1913). The combination of both extensions, logistic growth in the resource and a Holling type II functional response, results in the aforementioned Rosenzweig-MacArthur (RM) model (Rosenzweig and MacArthur 1963). The dynamics of the RM model are given by:

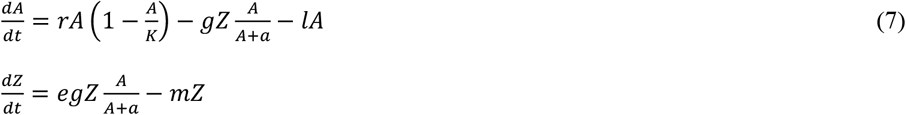

The expressions of the equilibria are (for Z^*^ > 0):

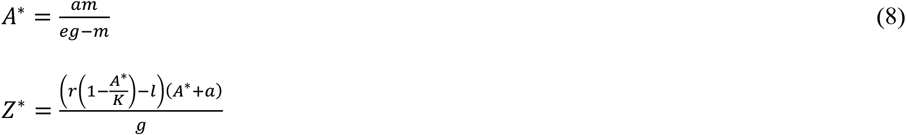

The Holling type II functional response term is part of both the equation of the resource and the consumer and therefore affects all elements of the Jacobian matrix. The partial derivative of the functional response term is 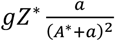. The expressions of the elements of the Jacobian Matrix are:

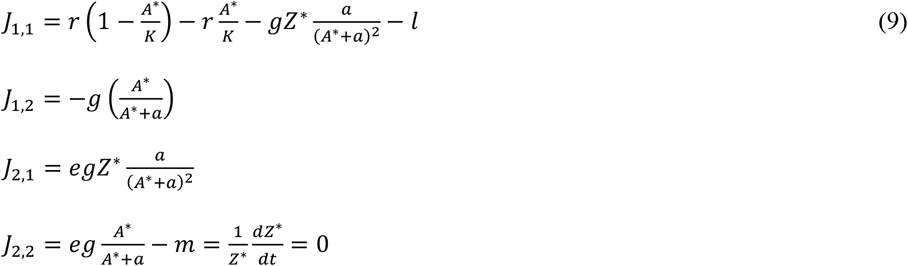

#### The Rosenzweig-MacArthur-Scheffer model

A third key extension of the LV model stems from the notion that above a given threshold density the consumer itself might become an attractive resource for a top-consumer. Such prey switching responses in the top-consumer can be modelled with a sigmoid Holling type III functional response of the top-consumer, which is a specific case of a Hill function (Hill 1910). A RM model with a Hill function in the loss term of the consumer was analyzed by Scheffer et al. (2000). We will further refer to this model as the Rosenzweig-MacArthur-Scheffer (RMS) model. In the RMS model a nonlinear Holling type III loss term of the consumer 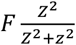 is added to the linear loss term of the LV model *mZ*. To fully comply with the formulation used by Scheffer et al. (2000) a constant influx of the resource is also added, thus mimicking chemostat dynamics. The dynamics of the RMS model are given by:

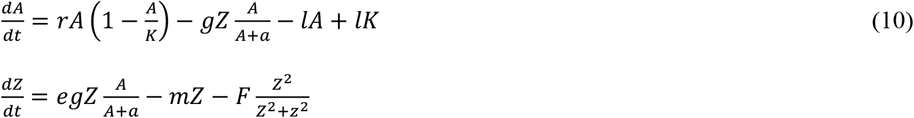

For the RMS model there are no explicit expressions of the equilibria, which need to be solved numerically. The addition of the nonlinear Holling type III in the consumer only affects the element *J*_*2.2*_ (compare Eq. 9). The partial derivative of the functional response is 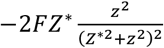. The expressions of *J*_*2.2*_ is:

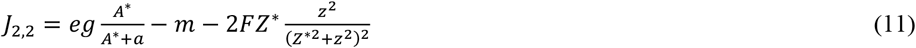

Each of these models is mathematically fully analyzed through bifurcation analysis for their stability properties and described in the literature (e.g. Rosenzweig and MacArthur 1963; Scheffer et al. 2000; Kooi 2003). The stability properties range from neutrally stable (LV), always stable (LVV), for some parameter values stable, for others unstable (RM) and for some parameter values showing alternative stable states (RMS).

### The linearization of functional response terms

In our linearization step, we replace the nonlinear growth-, interaction- and loss terms in the LVV, RM and RMS models by a single parameter. To maintain a link with the original models, and to allow an analysis of the linearized models along the same environmental gradients against which the original models can be analyzed, we express these linear parameters in terms of the original parameters *r, K, g, a, F* and *z*, and the equilibrium densities A* and Z*. By doing so, we guarantee that the equilibrium biomass densities and material flows of the original LVV, RM and RMS models are by definition equal to those for their linearized counterparts. Hence, we are able to analyze the same equilibria but using Lotka-Volterra dynamics instead of the original non-linear dynamics. We identify the linearized models with a reference to the LV model: LVV_(LV)_ for the linearized Lotka-Volterra-Verhulst model, RM_(LV)_ for the linearized Rosenzweig-MacArthur model and RMS_(LV)_ for the linearized Rosenzweig-MacArthur-Scheffer model. For each of the linearized models we also present the explicit expressions of the elements of the Jacobian matrix. We first present the elements in terms of the new parameters *r*_*lin*_, *g*_*lin*_ and *F*_*lin*_ and thereafter reformulated each element of that matrix in terms of the original parameters *r, K, g, a, F* and *z*. This allows for a one to one comparison of the original Jacobian matrix with the Jacobian matrix of the linearized models.

#### The linearized Lotka-Volterra-Verhulst model LVV_(LV)_

The only parameter of the LVV model that is linearized is the resource growth rate *r*_*lin*_ (Eq 13) and this parameter increases with increasing carrying capacity *K*. The dynamics of are given by:

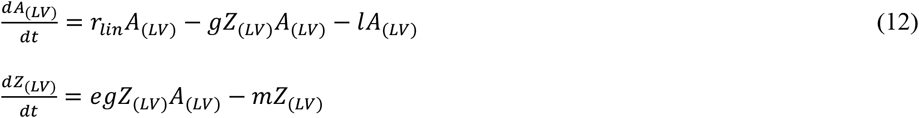

with

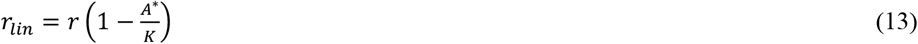

and:

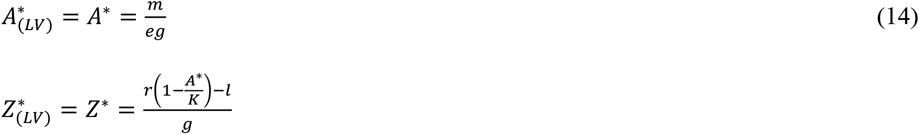

The expression of element *J*_*(LV)1.1*_ is (compare with Eq 6):

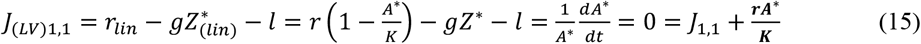

#### The linearized Rosenzweig-MacArthur model RM_(LV)_

During linearization of the RM model, two parameters are linearized, namely the resource growth rate *r*_*lin*_ and the interaction term *g_lin_* (Eq 17). However, of this two, only the resource growth rate increases with increasing carrying capacity *K*. The interaction term *g*_*lin*_ stays constant because its only variable is the resource concentration, which itself does not depend on the carrying capacity. The dynamics are given by:

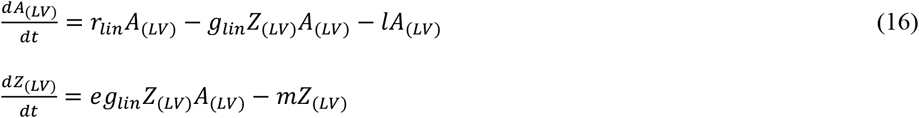

with

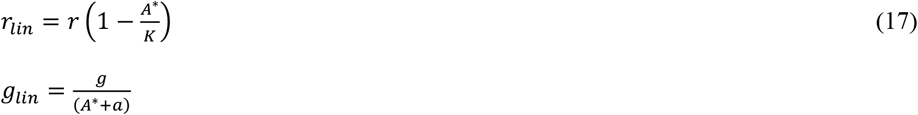

The expressions of the equilibria of linearized RM model are:

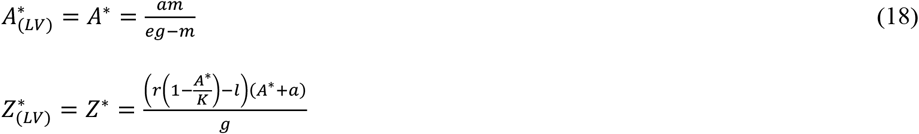

After reformulation, the partial derivative of the linearized Holling type II functional response is expressed as 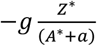. The full expressions of the elements of the linearized Jacobian matrix are (compare with Eqs 9):

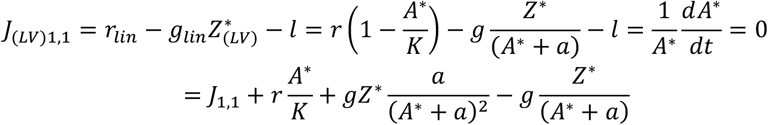

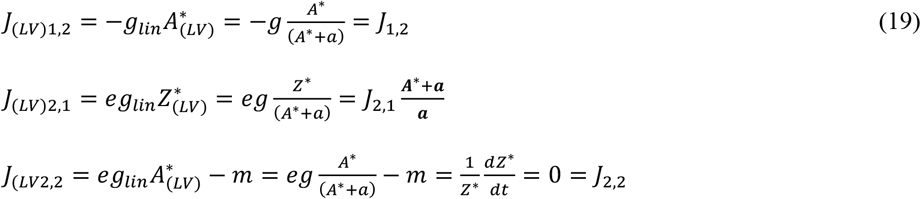

#### The linearized Rosenzweig-MacArthur-Scheffer model RMS_(LV)_

During linearization of the RMS model, three parameters are linearized, namely the resource growth rate *r*_*lin*_, the interaction term *g*_*lin*_ and the consumption rate by the top-consumer *F*_*lin*_ (Eq 21). Each of these linearized parameters changes with increasing top-consumer consumption rate or carrying capacity. The dynamics are given by:

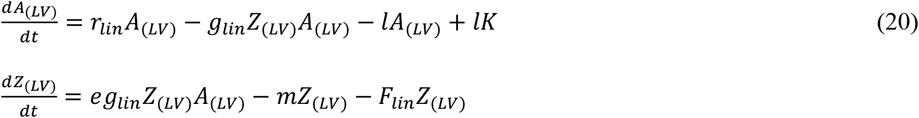

with:

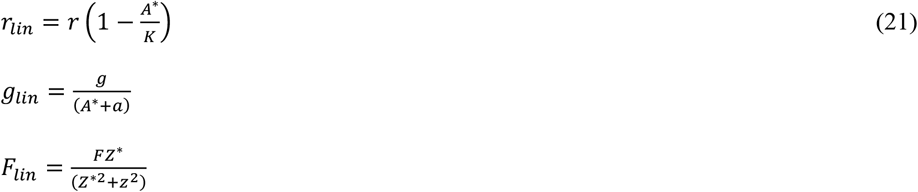

and:

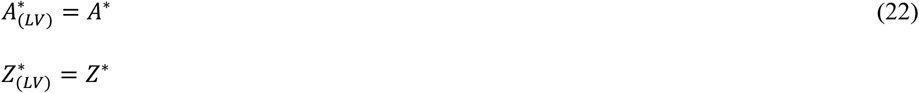

After reformulation, the partial derivative of the Holling type III functional response term is expressed as: 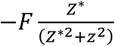. The full expression of element *J*_*(LV)2.2*_ is (compare with Eq 11):

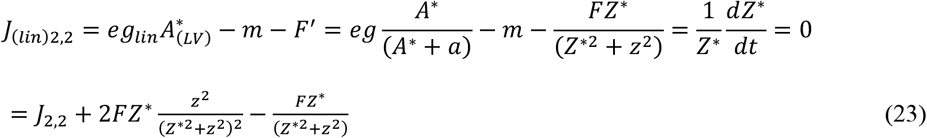

By replacing all non-linear terms with linear terms we imposed LV dynamics on the LVV, RM and RMS systems. However, we acknowledge that many empirical food web models presented in the literature actually include self-limitation (Moore and de Ruiter 2012; Van Altena et al. 2014; Dakos 2018). Therefore, we additionally present partially linearized versions of the RM and the RMS models whereby the logistic growth term in the resource is maintained. Thus, instead of LV dynamics we impose LVV dynamics on the RM and the RMS systems. An added advantage of creating intermediate versions is that we obtain a more complete understanding of the contribution of the different non-linear terms. We identify these partially linearized models with reference to the LVV model: RM_(LVV)_ for the Rosenzweig-MacArthur model where the Holling type II functional response in the predation term is replaced by a Holling type I without ceiling, but the logistic growth in the resource is maintained, and RMS_(LVV)_ for the Rosenzweig-MacArthur-Scheffer model where the Holling type II and type III terms are linearized but the logistic growth in the resource is maintained. Finally, we also consider the RMS version where the Type III functional response term is linearized but the logistic growth term and the Holling type II functional response term are maintained, and identify this model as RMS_(RM)_. A complete overview of all equations, original and linearized, of the LV, LVV, RM and RMS models is presented in S1 Appendix.

### Stability analyses of the equilibria along an environmental gradient

The LVV, the RM and the RMS models contain the carrying capacity *K* of the resource and can therefore be analyzed along an environmental gradient representing eutrophication. The RMS model however is intended for analysis along another gradient, namely that of consumption of the consumer by a top-consumer expressed in parameter *F*. For easy comparison with the RMS model as it is described in literature, we therefore choose to present the analysis for *F* in the main results, and present the analysis for *K* in S2 Appendix.

We parametrized the models and calculated the equilibrium values along the environmental gradients. We used the same parameters values as were used by Scheffer et al. (2000), to simplify comparison with previous research (Table 1). Their parameters were inspired by algae-zooplankton dynamics. Next, we took the equilibrium solutions, as if they were observations sampled from the virtual reality of the models with nonlinear terms, and used them to parametrize the models with the linear terms. Both the linear and the nonlinear model versions thus describe the same equilibrium biomass values and material flows. This step mimics the development of empirical food web models to describe observed equilibria in ecosystems as described by De Ruiter et al. (1993, 1995). We used the elements of the Jacobian matrix to calculate the eigenvalues and characterize the dynamic behavior of the systems. To evaluate and compare the stability properties of both the non-linear and the linear models along the same environmental gradients, we present the equilibrium solutions, the real part of the dominant eigenvalues (Re(*λ*)) and the dynamic behavior of each system. The Re(*λ*) gives an indication of the rate of recovery of the system after a small perturbation and is used as a measure of engineering resilience. Additionally, we present phase planes and nullclines for all models at two points along the environmental gradient to provide more insight into system behavior.

## RESULTS

### Lotka-Volterra-Verhulst model

The LVV model goes through a predator invasion threshold (or transcritical bifurcation) at *K* = 0.64 and is stable for higher carrying capacities, first as a stable node and thereafter as a stable focus (Fig 2b and 3a, c). Above the predator invasion threshold the equilibrium density of the resource is not dependent on the carrying capacity because the loss rate of the consumers does not depend on *K* and therefore its *R*\* (cf. (Tilman 1982)) is constant over the environmental gradient (Fig 2a, line *A*\*). Instead, the increasing productivity of the system ends up in increasing biomass of the consumer (Fig 2a, line *Z*\*).

**Figure 2.**
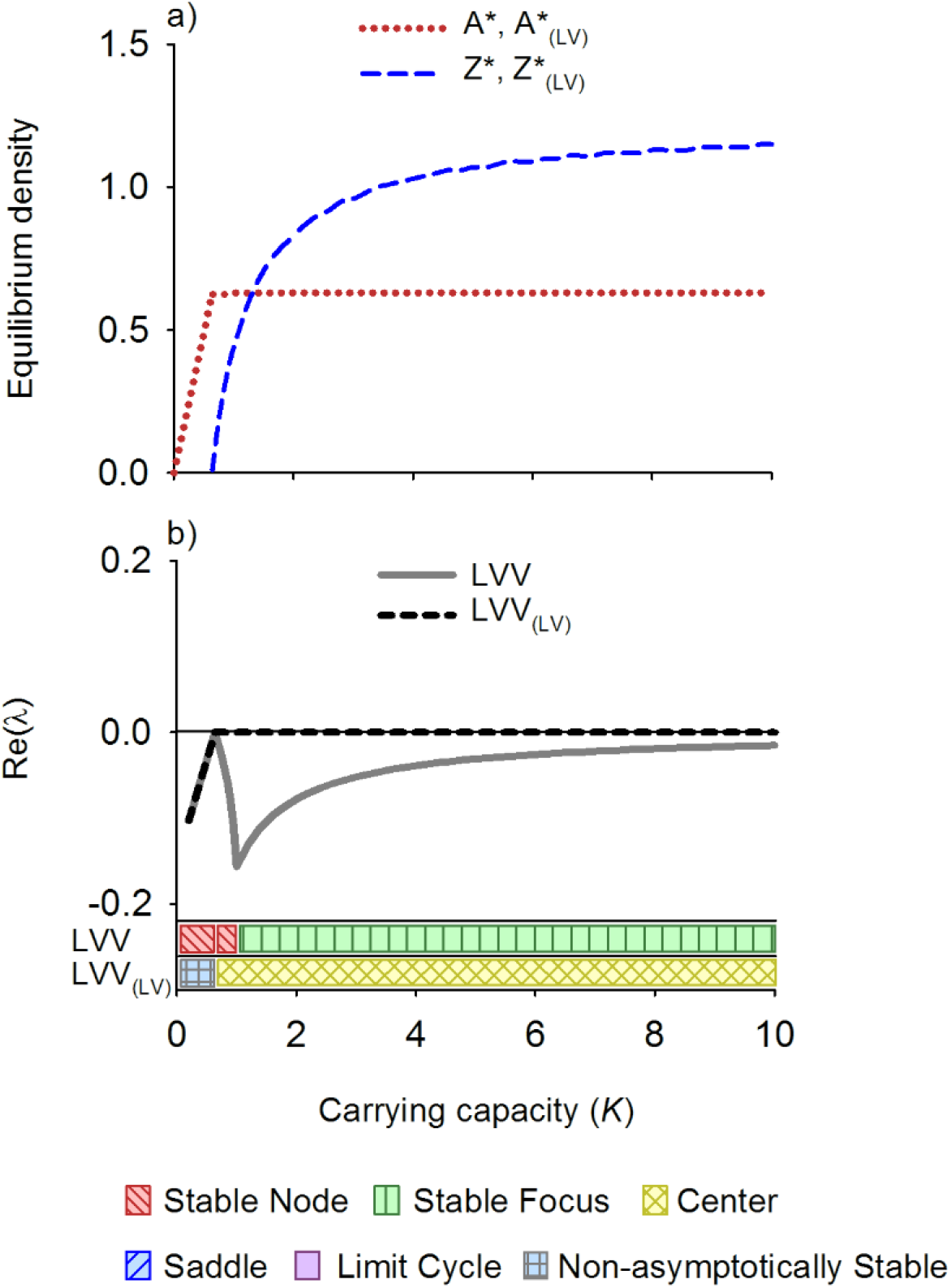
Equilibrium densities (**a**) and real part of the dominant eigenvalue (Re(*λ*)) (**b**) for the LVV model (Lotka-Volterra-Verhulst) and its counterpart with linear interaction terms, the LVV_(LV)_ model, along an environmental gradient. Also the dynamical behavior along the gradient is depicted, following Kot (2001).

**Figure 3.**
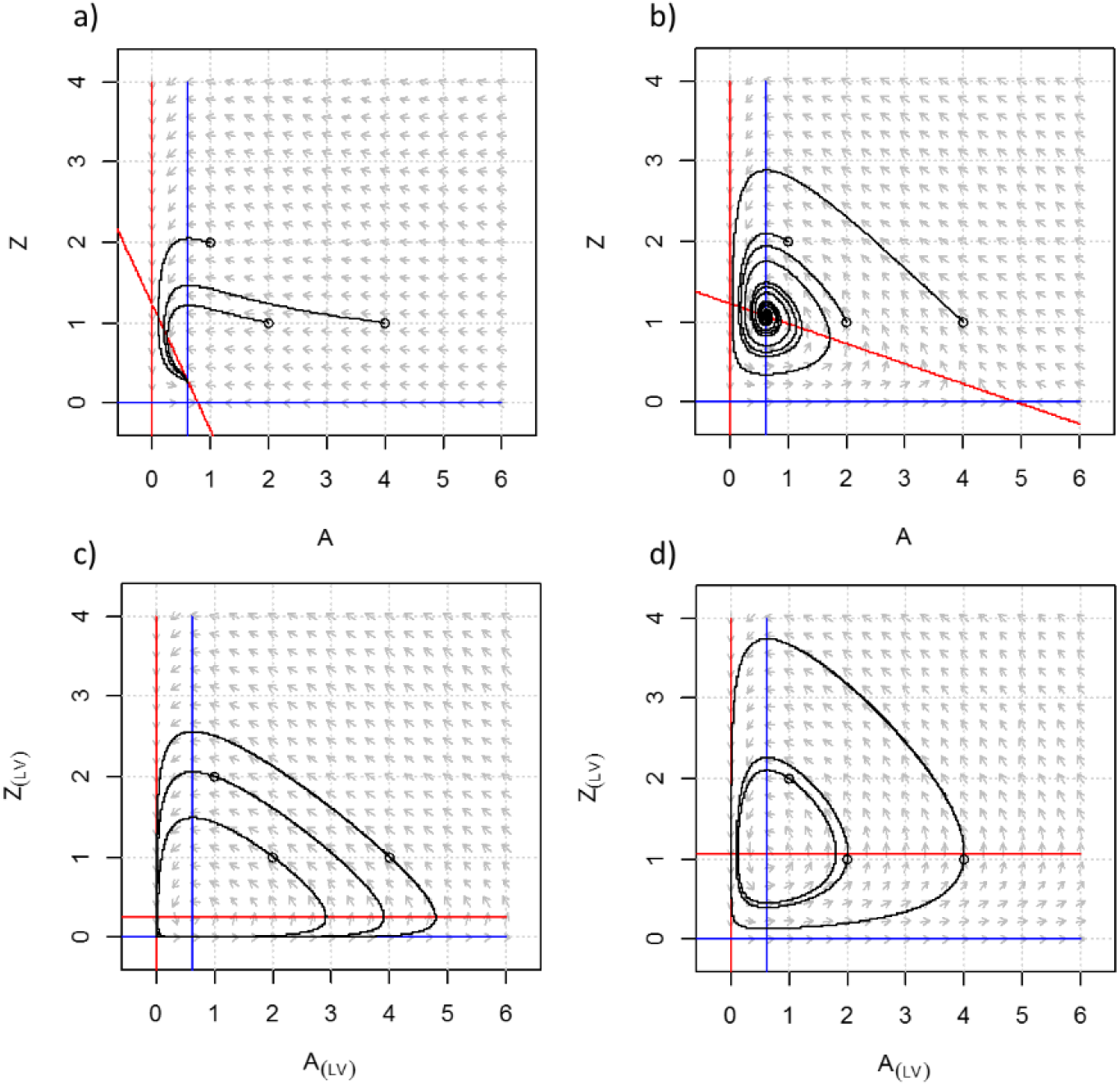
Phase planes and nullclines of the LVV model (Lotka-Volterra-Verhulst) for carrying capacity *K* = 0.8 (**a**) and *K* = 5 (**b**), and its counterpart with linear interaction terms, the LVV_(LV)_ model, for *K* = 0.8 (**c**) and *K* = 5 (**d**). While for each *K* the equilibrium solutions of the LVV and the LVV_(LV)_ systems are the same, the dynamical behavior differs distinctly. LVV shows a stable node (**a**) and a stable focus (**b**) while the LVV_(LV)_ is neutrally stable (**c-d**)

The linearized version of the LVV model, denoted as LVV_(LV)_, also shows a transcritical bifurcation at a corresponding value of *K* = 0.64 but thereafter shows the type of neutral stability that is typical for LV models with only linear terms (Fig 2b and 3b,d).

### Rosenzweig-MacArthur model

The RM model goes through a predator invasion threshold at *K* = 1.02, and shows a stable equilibrium thereafter, first as a stable node and thereafter as a stable focus (Fig 4a and 5a). At a value of *K* = 2.65 a supercritical Hopf bifurcation occurs and the model shows stable limit cycles as its dynamic behavior at higher carrying capacities (Fig 4a and 5b). This is the famous ‘paradox of enrichment’ that states that increasing the carrying capacity of the resource tends to destabilize consumer-resource interactions (Rosenzweig 1971). Also in the RM model, the equilibrium density of the resource is not dependent on the carrying capacity above the transcritical bifurcation because the loss rate of the consumer does not depend on *K* and therefore the *R*\* is constant over the environmental gradient (Fig 4a, line *A*\*). And again, the increasing productivity of the system ends up in increasing biomass of the consumer (Fig 4a, line *Z*\*).

The fully linearized version of the RM model, denoted as RM_(LV)_, also shows a transcritical bifurcation at the corresponding value of *K* = 1.02 and thereafter the type of neutral stability that is typical for LV models with only linear terms (Fig 4b and 5e,f). The intermediate version RM_(LVV)_ however, with logistic growth in the resource but a linear predation term, is stable after the critical invasion threshold, first as a stable node and thereafter as a stable focus (5c,d). Unlike the original RM, the RM_(LVV)_ does not show a supercritical Hopf bifurcation with increasing carrying capacity.

**Figure 4.**
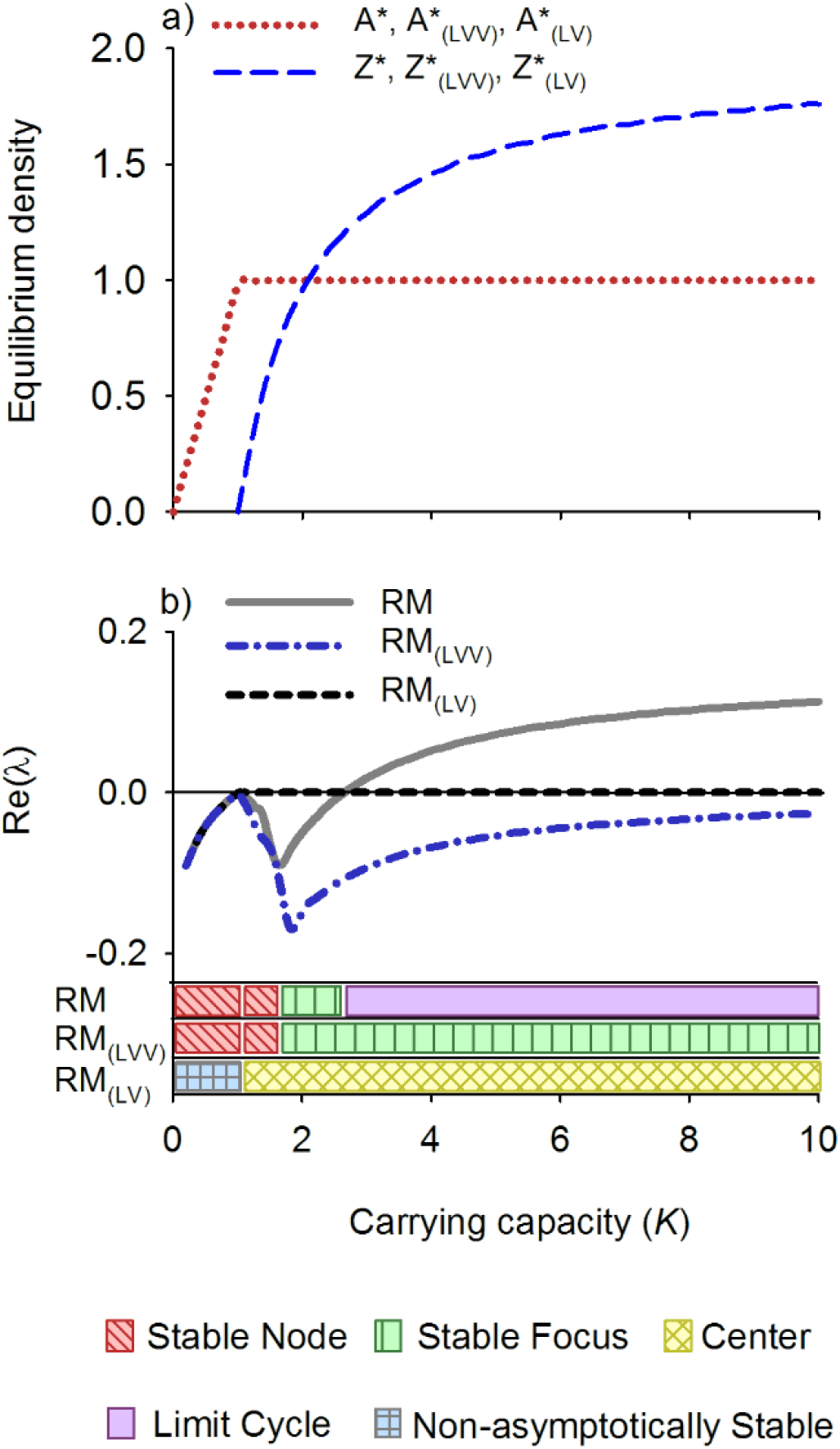
Equilibrium densities (**a**) and real part of the dominant eigenvalue (Re(*λ*)) (**b**) for the RM model (Rosenzweig-MacArthur), and its counterparts with linearized interaction terms, the RM_(LVV)_ and RM_(LV)_ models, along an environmental gradient. Also the dynamical behavior along the gradient is depicted, following Kot (2001).

**Figure 5.**
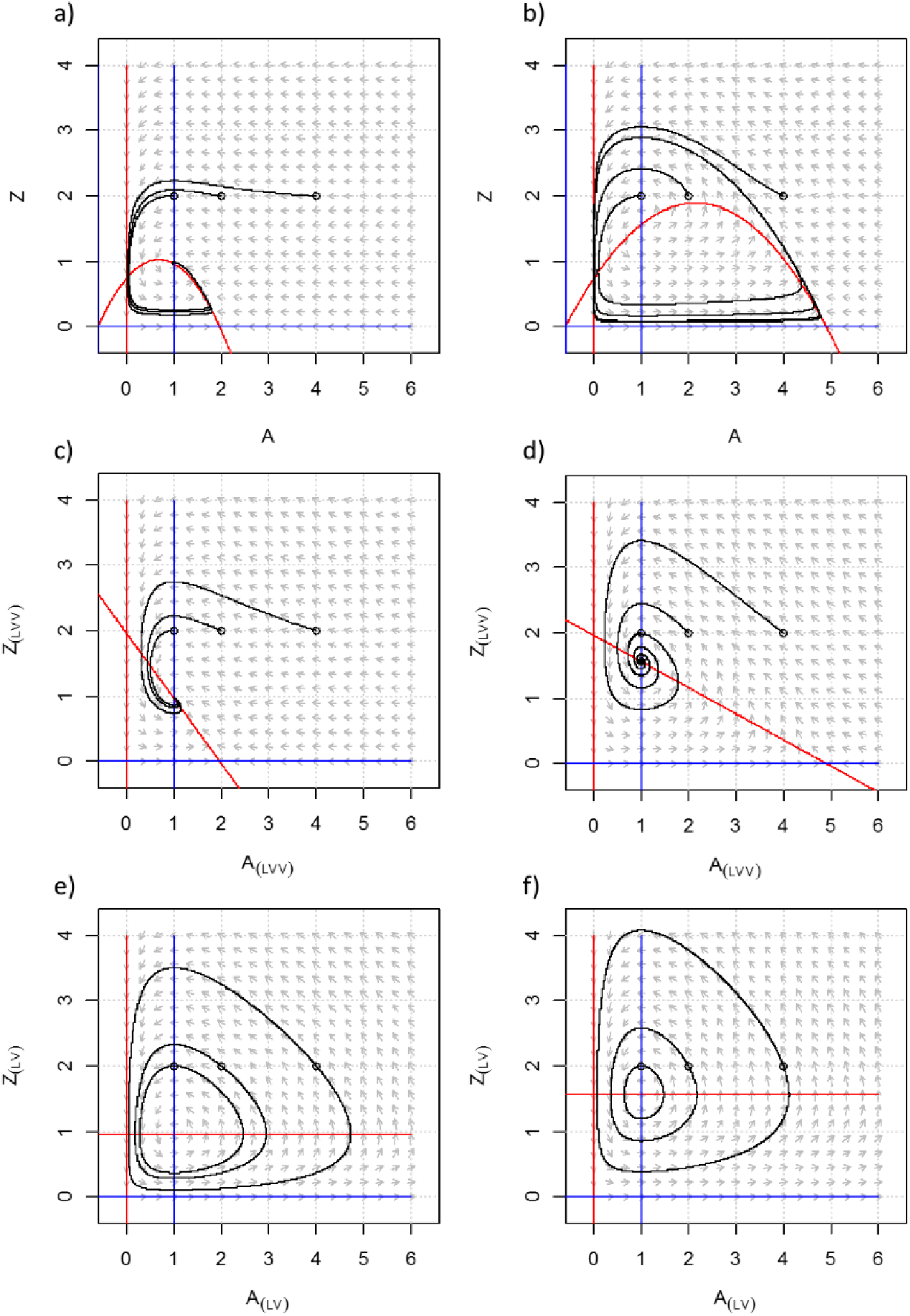
Phase planes and nullclines of the RM model (Rosenzweig-MacArthur) for carrying capacity *K* = 0.8 (**a**) and *K* = 5 (**b**), and its counterparts with linear interaction terms, the RM_(LVV)_ model for *K* = 2 (**c**) and *K* = 5 (**d**), and the RM_(LV)_ model for *K* = 2 (**e**) and *K* = 5 (**f**). While for each *K* the equilibrium solutions of the RM, RM_(LVV)_ and the RM_(LV)_ systems are the same, the dynamical behavior differs distinctly. RM shows a stable node (**a**) and a limit cycle (**b**), the RM_(LVV)_ only a stable node (**c-d**) while the RM_(LV)_ is neutrally stable (**e-f**)

### Rosenzweig-MacArthur-Scheffer model

The equilibrium density of both the resource and the consumer is dependent on the consumption rate of the top-consumer *F* (Fig 6a, line A*) and the model shows a complex response to changing top-consumer consumption rate. Namely, the RMS model shows alternative stable states with critical transitions at *F* = 0.2408 and *F* = 0.076 (Fig 6b). In addition to these saddle node bifurcations, the system shows a supercritical Hopf bifurcation at *F* = 0.2404. As a result, the system shows limit cycles with increasing amplitude at lower values of *F* (Fig 7a). When the amplitude becomes large enough, the system will shift to the other stable state via a homoclinic (global) bifurcation (Fig 6b and 7b). For more details on this aspect of the model see Scheffer et al. (2000). The results for enrichment (*K*) are discussed in S2 Appendix.

**Figure 6.**
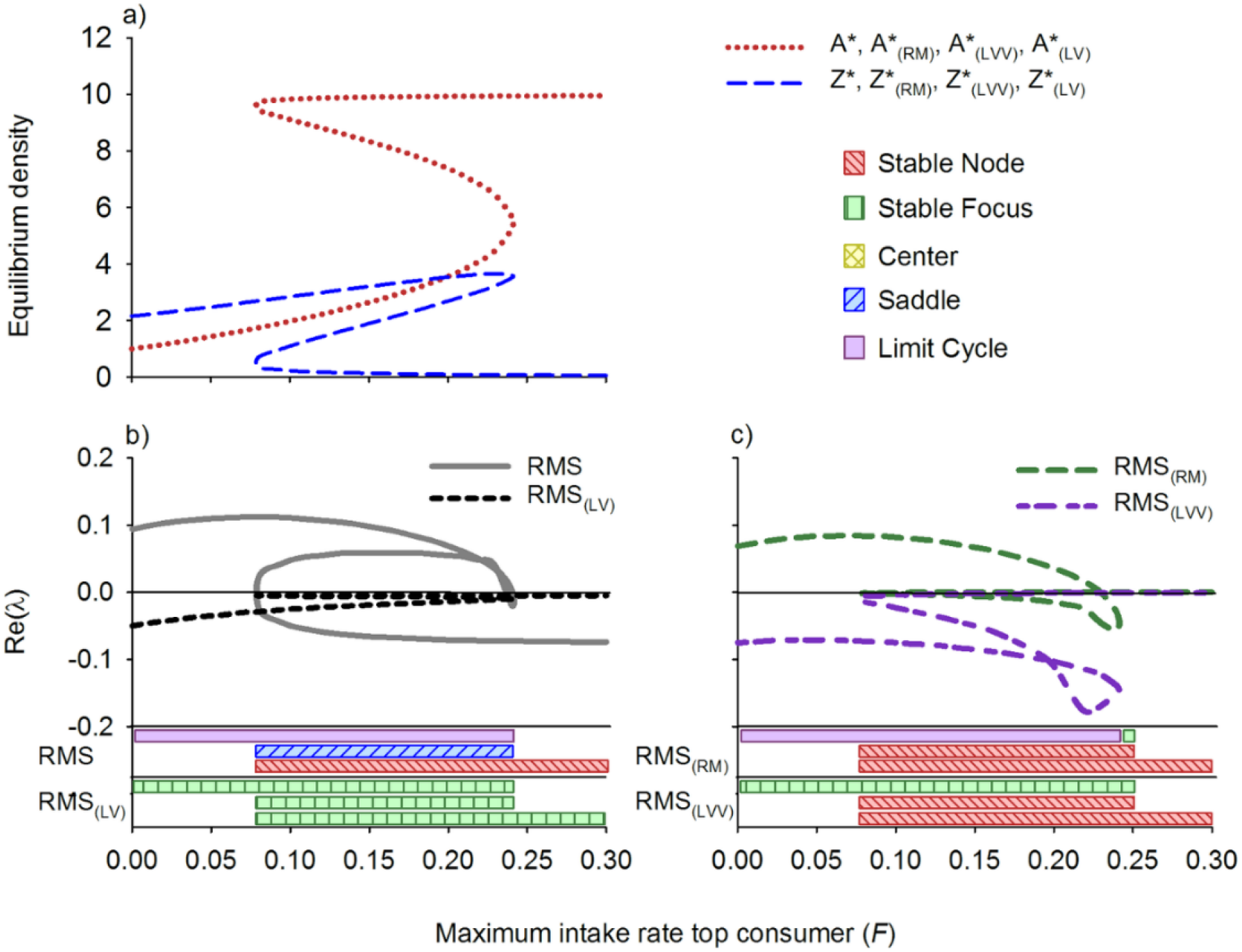
Equilibrium densities (**a**) for the RMS model (Rosenzweig-MacArthur) and its counterparts with linear interaction terms, the RMS_(lv)_ model, the RMS_(RM)_ model and the RMS_(LVV)_ model, and the real part of the dominant eigenvalue (Re(*λ*)) for (**b**) RMS and RMS_(lv),_ and (**c**) for RMS_(RM)_ and RMS_(LVV)_ along an environmental gradient. Also the dynamical behavior along the gradient is depicted, following Kot (2001). The carrying capacity *K* = 10.

**Figure 7.**
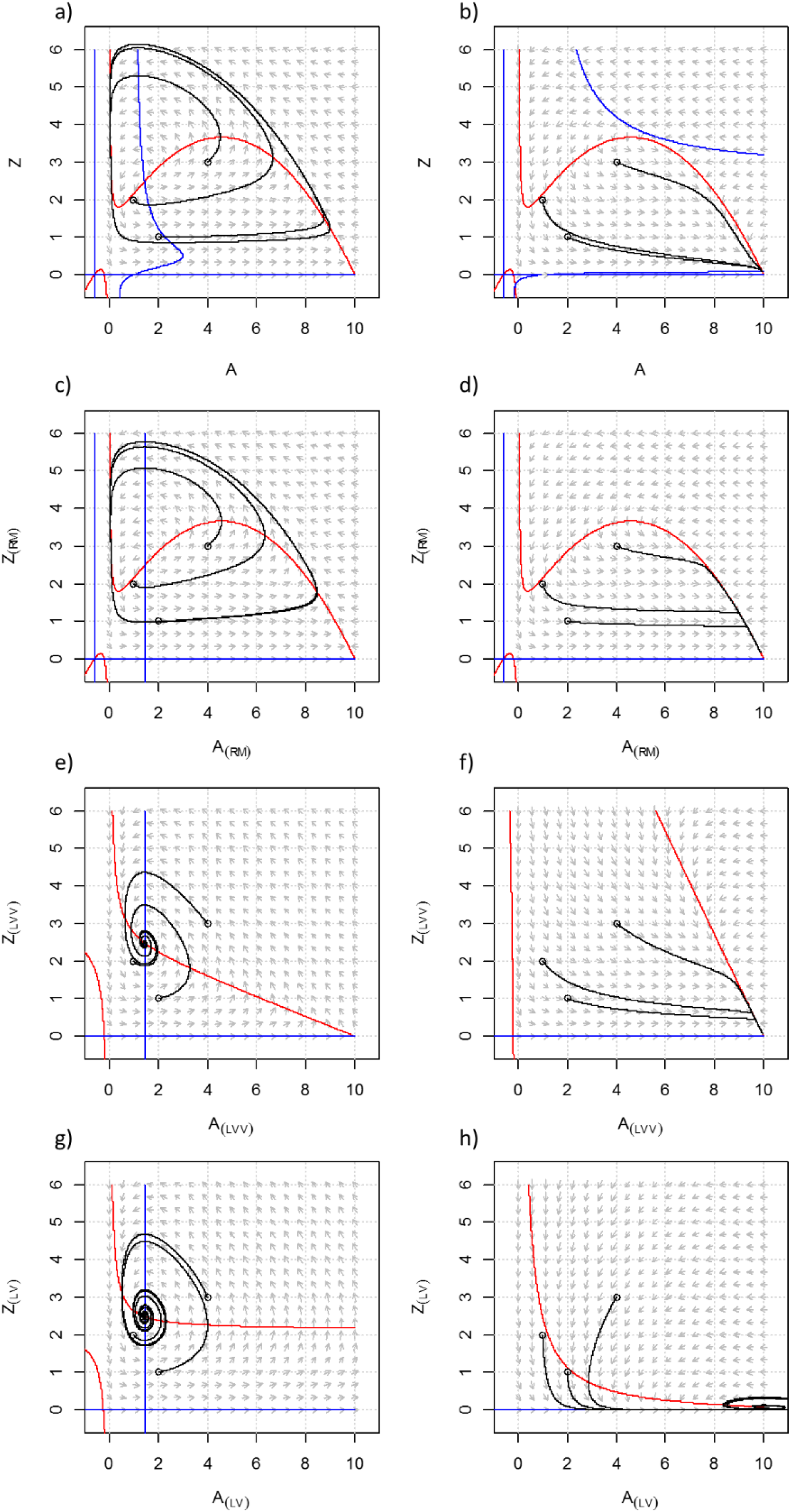
Phase planes and nullclines of the RMS model (Rosenzweig-MacArthur-Scheffer) for carrying capacity *F* = 0.05 (**a**) and *F* = 0.25 (**b**), and its counterparts with linear interaction terms, the RMS_(RM)_ model for *F* = 0.05 (**c**) and *F* = 0.25 (**d**), the RM_(LVV)_ model for *F* = 0.05 (**e**) and *F* = 0.25 (**f**) and the RM_(LV)_ model for *F* = 0.05 (**g**) and *F* = 0.25 (**h**). While for each value of *F* the equilibrium solutions of the RMS, RMS_(RM)_, RMS_(LVV)_ and RMS_(LV)_ systems are identical, their dynamical behavior differs distinctly. RMS and RMS_(RM)_ show a limit cycle (**a,c**) and a stable node (**b,d**) while the RMS_(LVV)_ shows stable focus and a stable node and the RMS_(LV)_ a stable focus (**b-d**)

The fully linearized version of the RMS model (denoted as RMS_(LV)_) shows stable focus for any value of top-consumer abundance (Fig 6b, 7g,h). This different behavior of the RMS_(LV)_ model compared with the LVV_(LV)_ and the MS_(LV)_ model can be explained from the chemostat dynamics that were built in the resource equation of the RMS model. Importantly, the RMS_(LV)_ model does not show (saddle node) bifurcations. Nonetheless, the Re(*λ*) of RMS_(LV)_ appears to incrementally decrease in the direction of the bifurcation point at *F* = 0.2408 in the RMS model (Fig 6b). For the bifurcation point in the opposite direction however there is no trace of such pattern. The RMS model with RM dynamics (denoted as RMS(RM)) first shows limit cycles, similar to the original RMS model (Fig 6c, 7c). However, the limit cycles turn into stable focus through a Hopf bifurcation and subsequently into a stable node (Fig 6c, 7d). The saddle node bifurcation has disappeared. The RMS model with LVV dynamics (denoted as RMS(LVV)) shows no bifurcations (Fig 6c). The equilibria are stable for all values of *F*, either as a stable focus (Fig 7e) or stable node (Fig 7f).

## DISCUSSION

When models are constructed to describe the dynamics of ecosystems, key choices have to be made about which processes to include and how these processes are formulated. These choices are typically shaped by the goal of the modelling project (IPBES 2016), but a complicating factor is that models with different levels of detail and complexity can be developed to accurately describe the same empirical information such as biomass densities and feeding rates. A principle challenge for theoretical ecologists is to understand the consequence of using alternative model formulations for capturing the stability of ecosystems, and how to translate between these formulations to create useful knowledge to understand and anticipate the impacts of ongoing global environmental change (Fulton et al. 2003a, b; Aldebert et al. 2016a, 2018). The classical sensitivity analysis quantifies how changes of environmental or physiological model parameters propagate into an uncertainty in model predictions. A fairly recent extension is structural sensitivity analysis where the consequences of changes of whole building blocks of the model are studied (see Aldebert et al. 2016b and references therein). The approach we present in this study can also be classified as a structural sensitivity analysis of a number of classical consumer resource models. However, a key difference is that the equilibrium solutions of our models are kept identical, so that we can focus on how the analysis of the local stability of the equilibrium solutions depends on the use of alternative model formulations for describing the equilibria (Flora et al. 2011; Adamson and Morozov 2013).

Our main finding is that changing nonlinear terms in minimal dynamic models of consumer-resource interactions into linear ones, while maintaining the equilibrium densities and hence the flux of mass, has a strong impact on the stability properties of models, reducing the range of dynamics they can exhibit and decreasing the number of variety of reorganizations they can experience (Abrams 2001; Adamson and Morozov 2013). The transcritical bifurcation in the RM model, that is, the transition from an equilibrium of a very small population size to extinction, is captured by the linearized RM_(LVV)_ model, and partly by the RM_(LV)_ model (Fig 4b). However, linearized models fail to consistently capture the Hopf bifurcations, when the system becomes cyclic, or saddle-node bifurcations, when the system undergoes a critical regime shift (Fig 4b, 6b,c). Thus, the eigenvalues of the simpler model systems with only linear interaction terms do not provide an indication of the ecological resilience of the original non-linear model systems.

Based on the direct comparison of the analytical expressions of the Jacobian elements of the original and linearized models (Eq 15, 19, 23) we could already expect that the resulting matrices would show different and non-coupled equilibrium dynamics along the environmental gradient, despite their identical equilibrium solutions. Indeed, some may label the findings as presented here as somewhat trivial considering that it has long been known that non-linear interaction terms can have drastic effects on the stability of dynamic systems. We, however, argue that these results represent an important basis against which the scope, outcomes and limitations of local stability analyses of real-world ecosystem need to be discussed (Fig 1). The theoretical experiments performed here may be compared with the situation where empirical food web models with only linear interspecific interaction terms are used to analyze the stability of real-world ecosystems that are inherently complex and nonlinear (Fig. 1; Abrams 2001; Moore and de Ruiter 2012). Our findings suggest that we should be cautious when conferring the stability properties of empirical food web models that assume linear interaction terms to the actual ecosystem under study, given that the stability properties of the modelled equilibria appeared to be controlled by the assumptions on underlying dynamics rather than the observed equilibrium solutions (Gross et al. 2004). This notion is especially relevant in light of ecological resilience and environmental change, as our results suggest that insights from stability analyses of observed food webs cannot simply be extrapolated into the realm of regime shifts (Fig. 1).

The failure of linearized models to consistently signal regime shifts in our analysis contrasts with the results found by Kuiper et al. (2015) who used the complex ecosystem model PCLake (Janse 1997; Janse et al. 2010) as a virtual reality in which they sampled information to parameterize ‘empirical’ food web models consisting of only linear interaction terms. They found that the stability of the food web models decreased towards both the regime shift from clear- to turbid-water during eutrophication and one from turbid- to clear-water during re-oligotrophication. Importantly, however, PCLake contains a more realistic food web description compared to the minimal dynamical models used in this study. Apparent non-random patterns in network topology, such as connectence (Van Altena et al. 2016), feedback loops (Neutel et al. 2002) and the distribution of weak links (Emmerson and Yearsley 2004) are present in both PCLake and the linearized descriptions of tis food web. Apparently, the organization of biomasses and energy flows contains enough information to reveal the actual stability properties of the system. This is consistent with the dominant view among food web ecologists that stability is woven by non-random structures in complex webs (Polis 1998; Moore and de Ruiter 2012), and the rationale for developing food web models in the first place. Yet, by showing here that for simpler models the type of the functional response is decisive, the question arises how many realistic food web patterns should be represented in food web models to ensure that the calculated stability matric is not an artifact. We see complementary paths forward to answer this question.

The first is to better validate ecological models at the level of the emergent stability properties, by comparing the stability properties of models with the stability properties of the real ecosystems that are portrayed. There has been great progress in the development of empirical techniques for identifying and quantifying trophic interactions for the development of empirical food web models (Heijboer et al. 2017; Rosenbaum and Rall 2018). Yet the emergent stability properties of empirical food web models, resulting from the interplay of all the interactions, and as calculated by the Jacobian matrix approach, have rarely been tested against real world dynamics. An obvious reason is that system-wide experimental perturbations and monitoring actions are costly and often unfeasible. Fortunately, there is increasing knowledge about how local experimental or natural perturbations may be used to reveal system-wide ecological resilience, and which species serve best as indicators (Kuiper et al. 2015; Van de Leemput et al. 2017; Dakos 2018). For example, in shallow lakes, the regrowth rate of aquatic vegetation after harvesting a fraction of the biomass may indicate system-wide ecological resilience (Kuiper et al. 2017; Van de Leemput et al. 2017). Furthermore, we see much potential in the continuation of the studies where series of empirical food web models are established along environmental gradients in real ecosystems (Neutel et al. 2007; Andres et al. 2016; Schwarz et al. 2017). Such analyses could be repeated for ecosystems that are known to show abrupt regime shifts, like shallow lakes and peatland ecosystems (Rocha et al. 2015), for example by performing mesocosm experiments (Moss et al. 2004) or by making paleoecological reconstructions of food webs (Rawcliffe et al. 2010).

An complementary way forward may be to replace the linear functional response terms of empirical food web models into more realistic non-linear terms (Kalinkat et al. 2013), thus creating models that are complex both in terms of the number of equations and the shape of the functional response used to characterize interactions. There are numerous forms of hybrid models presented in the literature, such as extended minimal models that represent highly simplified food webs (e.g. McCann and others 1998; Kooi 2003; Rooney and others 2006) and theoretical models of large webs with non-linear interaction (e.g. Gross and others 2009). An important reason for the current use of linear terms is a lack of available information to parameterize more sophisticated functions (Neutel and Thorne 2016). Yet recent progress in development of methods for quantifying trophic interactions (Heijboer et al. 2017) and sharing of data (Culina et al. 2018) will take away many of the existing limitations. Especially Holling type II functional response terms are commonly used to describe consumer-resource interactions and compelling examples exist where functional response terms have been accurately parametrized with experimental data (Sentis et al. 2012; Kalinkat et al. 2013). Neutel and Thorne (2016) conducted a structural sensitivity analysis of an empirically derived food web model of an Antarctic dry tundra ecosystem, replacing type I functional response terms with type II and type III. They concluded that unless populations receive their main energy input from external sources, such as in lakes, it is implausible that saturation effects in predation play a significant role in multi-trophic equilibrium dynamics. However, the implications of solely using type I functional response terms for modelling and assessing the ecological resilience of terrestrial Antarctic ecosystems, that is, detecting decreasing resilience towards a regime shift in response to environmental change, remain mostly unexplained.

Arguably, the local stability analysis based on the Jacobian matrix approach does not provide a complete framework for studying ecosystem long-term dynamics analysis in the context of environmental change. Complementary approaches that focus on transients, for example, have been proposed (Neubert and Caswell 1997; Hastings 2004). Alternatively, large simulation models may be expended to better connect ecosystem dynamics with food web theory. The food web structure in simulation models is often reduced to its bare essentials, to balance complexity and optimize performance (Fulton et al. 2003b). It is therefore that Hannah et al (2010) call for a more comprehensive description of the structure of food webs in the designs of the next generation ecosystem models. An evident downside of increasing model complexity is that it goes at the costs of mathematical tractability, and hence, scientific understanding (Scheffer and Beets 1994). Solutions may be found in increases of data availability, computational resources, and new approaches to modelling such machine learning (Purves et al. 2013). At the same time, the combination of data-spare regions, shifting conditions, and irreducible complexity suggests that the need for multiple modelling perspectives and their bridging will continue to increase.

Connecting modellers and other scientists who are seeking to understand how food webs shape and are reshaped by their interactions across space, abiotic processes, as well as by human action and structures, requires bridging across knowledge systems. A strategy to overcome the limitations of single modelling framework is to exploit the diversity of modelling approaches (*cf.* Janssen et al. 2015), for example by applying them side by side within one integrated environmental assessment (Logan 1994; Weijerman et al. 2015). Indeed, it has been postulated that better aligning, integrating and reconciling adjacent research fields is key to furthering science (e.g. Thompson et al. 2012; Barnes et al. 2018; Kadowaki et al. 2018). Recently, there has been a new interest in techniques for bridging different knowledge, especially in the context of science and practice, but such techniques are also extremely relevant for bridging different communities of practice within science (Tengö et al. 2014, 2017). The multiple evidence base approach of Tengö et al. (2014, 2017) proposes that bridging knowledge systems requires: *knowledge mobilization, translation, negotiation, application*, and *synthesis*. The mobilization of multiple types of knowledge is required to share knowledge in forms that others can understand; translation between knowledge systems is often required to enable mutual comprehension of shared knowledge; negotiation enables a joint assessment of convergence, divergence, and conflicts across knowledge contributions; synthesis concerns the shaping of broadly accepted common knowledge bases for a particular purpose, respecting the integrity of each knowledge system, and the application of knowledge enables the creation of outputs that are designed to be useful for understanding different types of ecosystems facing a variety different decision contexts (IPBES 2016). Our results highlight that there is a great need for a multiple evidence base approach in ecology, to consolidate the insights and tools produced by different modelling approaches, and obtain a coherent understanding of how the Biosphere responds to the human force.

## Supporting information

Full presentation of equations

## ACKNOWLEDGEMENTS

We thank Cassandra van Altena, Luuk P.A. van Gerven, Annette B.G. Janssen, Wobbie van den Hurk for their help with programming and the mathematics. We thank Jan H. Janse for his valuable comments on the manuscript. JK received support from the Netherlands Foundation for Applied Water Research (STOWA) Project No. 443237, the Swedish Research Counsel FORMAS Project No. 2018-02371 and the Marianne and Marcus Wallenberg Foundation Research Exchange Program on Natural Capital, Resilience and Biosphere Stewardship.

