## Supplementary material for "Bridging theories for ecosystem stability through structural sensitivity analysis of ecological models in equilibrium": Full presentation of equations

### S1 Appendix

#### The Lotka-Volterra model - LV

The dynamics of the LV model are given by:

$$\frac{dA}{dt} = rA - gZA - lA \qquad \text{(S Eq. 1)}$$

$$\frac{dZ}{dt} = egZA - mZ$$

The expressions of the equilibria are:

$$A^* = \frac{m}{eg} \qquad \text{(S Eq. 2)}$$

$$Z^* = \frac{r - l}{g}$$

The expressions of the elements of the Jacobian matrix are:

$$J_{1,1} = r - gZ^* - l = \frac{1}{A^*}\frac{dA^*}{dt} = 0 \qquad \text{(S Eq. 3)}$$

$$J_{1,2} = -gA^* = -g\frac{m}{eg} = \frac{m}{e}$$

$$J_{2,1} = egZ^* = eg\frac{r-l}{g} = e(r - l)$$

$$J_{2,2} = egA^* - m = \frac{1}{Z^*}\frac{dZ^*}{dt} = 0$$

#### The Lotka-Volterra-Verhulst model - LVV

The dynamics of the LVV model are given by:

$$\frac{dA}{dt} = rA\left(1 - \frac{A}{K}\right) - gZA - lA \qquad \text{(S Eq. 4)}$$

$$\frac{dZ}{dt} = egZA - mZ$$

The expressions of the equilibria are (for $Z^* > 0$):

$$A^* = \frac{m}{eg} \qquad \text{(S Eq. 5)}$$

$$Z^* = \frac{r\left(1-\frac{A^*}{K}\right)-l}{g}$$

The expressions of the elements of the Jacobian Matrix are:

$$J_{1,1} = r\left(1 - \frac{A^*}{K}\right) - r\frac{A^*}{K} - gZ^* - l \qquad \text{(S Eq. 6)}$$

$$J_{1,2} = -gA^*$$

$$J_{2,1} = egZ^*$$

$$J_{2,2} = egA^* - m = \frac{1}{Z^*}\frac{dZ^*}{dt} = 0$$

**The Lotka-Volterra-Verhulst model with LV dynamics - $\mathrm{LVV}_{(LV)}$**

The dynamics of the $\mathrm{LVV}_{(LV)}$ model are given by:

$$\frac{dA_{(LV)}}{dt} = r_{lin}A_{(LV)} - gZ_{(LV)}A_{(LV)} - lA_{(LV)} \qquad \text{(S Eq. 7)}$$

$$\frac{dZ_{(LV)}}{dt} = egZ_{(LV)}A_{(LV)} - mZ_{(LV)}$$

with

$$r_{lin} = r\left(1 - \frac{A^*}{K}\right) \qquad \text{(S Eq. 8)}$$

The expressions of the equilibria of the $\mathrm{LVV}_{(LV)}$ model are (for $Z^* > 0$):

$$A^*_{(LV)} = A^* = \frac{m}{eg} \qquad \text{(S Eq. 9)}$$

$$Z^*_{(LV)} = Z^* = \frac{r\left(1-\frac{A^*}{K}\right)-l}{g}$$

The expressions of the elements of the Jacobian matrix of the $\mathrm{LVV}_{(LV)}$ are (compare with S Eq. 6):

$$J_{(LV)1,1} = r_{lin} - gZ^*_{(LV)} - l = r\left(1 - \frac{A^*}{K}\right) - gZ^* - l = \frac{1}{A^*}\frac{dA^*}{dt} = 0 = J_{1,1} + \boldsymbol{\frac{rA^*}{K}} \qquad \text{(S Eq. 10)}$$

$$J_{(LV)1,2} = -gA^*_{(LV)} = -gA^* = J_{1,2}$$

$$J_{(LV)2,1} = egZ^*_{(LV)} = egZ^* = J_{2,1}$$

$$J_{(LV)2,2} = egA^*_{(LV)} - m = egA^* - m = \frac{1}{Z^*}\frac{dZ^*}{dt} = 0 = J_{2,2}$$

**The Rosenzweig-MacArthur model - RM**

The dynamics of the RM model are given by:

$$\frac{dA}{dt} = rA\left(1 - \frac{A}{K}\right) - gZ\frac{A}{A+a} - lA \qquad \text{(S Eq. 11)}$$

$$\frac{dZ}{dt} = egZ\frac{A}{A+a} - mZ$$

The expressions of the equilibria are (for $Z^* > 0$):

$$A^* = \frac{am}{eg-m} \qquad \text{(S Eq. 12)}$$

$$Z^* = \frac{\left(r\left(1-\frac{A^*}{K}\right)-l\right)(A^*+a)}{g}$$

The expressions of the elements of the Jacobian Matrix are:

$$J_{1,1} = r\left(1 - \frac{A^*}{K}\right) - r\frac{A^*}{K} - gZ^*\frac{a}{(A^*+a)^2} - l \qquad \text{(S Eq. 13)}$$

$$J_{1,2} = -g\left(\frac{A^*}{A^*+a}\right)$$

$$J_{2,1} = egZ^*\frac{a}{(A^*+a)^2}$$

$$J_{2,2} = eg\frac{A^*}{A^*+a} - m = \frac{1}{Z^*}\frac{dZ^*}{dt} = 0$$

**The Rosenzweig-MacArthur model with LVV dynamics – RM$_{(LVV)}$**

The dynamics of the RM$_{(LVV)}$ model are given by:

$$\frac{dA_{(LVV)}}{dt} = rA_{(LVV)}\left(1 - \frac{A_{(LVV)}}{K}\right) - g_{lin}Z_{(LVV)}A_{(LVV)} - lA_{(LVV)} \quad \text{(S Eq. 14)}$$

$$\frac{dZ_{(LVV)}}{dt} = eg_{lin}Z_{(LVV)}A_{(LVV)} - mZ_{(LVV)}$$

with

(S Eq. 15)

$$g_{lin} = \frac{g}{(A^* + a)}$$

The expressions of the equilibria of the RM' model are (for $Z^* > 0$):

$$A^*_{(LVV)} = A^* = \frac{am}{eg-m} \quad \text{(S Eq. 16)}$$

$$Z^*_{(LVV)} = Z^* = \frac{\left(r\left(1-\frac{A^*}{K}\right)-l\right)(A^*+a)}{g}$$

The expressions of the elements of the Jacobian matrix are (compare with S Eq. 13):

$$J_{(LVV)1,1} = r\left(1 - \frac{A^*_{(LVV)}}{K}\right) - r\frac{A^*_{(LVV)}}{K} - g_{lin}Z^*_{(LVV)} - l = r\left(1 - \frac{A^*}{K}\right) - r\frac{A^*}{K} - g\frac{Z^*}{(A^* + a)} - l$$

$$= J_{1,1} + gZ^*\frac{a}{(A^* + a)^2} - g\frac{Z^*}{(A^* + a)}$$

$$J_{(LVV)1,2} = -g_{lin}A^*_{(LVV)} = -g\frac{A^*}{(A^*+a)} = J_{1,2} \quad \text{(S Eq. 17)}$$

$$J_{(LVV)2,1} = eg_{lin}Z^*_{(LVV)} = eg\frac{Z^*}{(A^*+a)} = J_{2,1}\frac{\boldsymbol{A^*+a}}{\boldsymbol{a}}$$

$$J_{(LVV)2,2} = eg_{lin}A^*_{(LVV)} - m = eg\frac{A^*}{(A^*+a)} - m = \frac{1}{Z^*}\frac{dZ^*}{dt} = 0 = J_{2,2}$$

**The Rosenzweig-MacArthur model with LV dynamics - RM$_{(LV)}$**

The dynamics of the RM$_{(LV)}$ model are given by:

$$\frac{dA_{(LV)}}{dt} = r_{lin}A_{(LV)} - g_{lin}Z_{(LV)}A_{(LV)} - lA_{(LV)}$$

(S Eq. 18)

$$\frac{dZ_{(LV)}}{dt} = eg_{lin}Z_{(LV)}A_{(LV)} - mZ_{(LV)}$$

with

$$r_{lin} = r\left(1 - \frac{A^*}{K}\right) \quad \text{(S Eq. 19)}$$

$$g_{lin} = \frac{g}{(A^*+a)}$$

The expressions of the equilibria of the RM$_{(LV)}$ model are (for $Z^*_{(LV)} > 0$):

$$A^*_{(LV)} = A^* = \frac{am}{eg-m} \quad \text{(S Eq. 20)}$$

$$Z^*_{(LV)} = Z^* = \frac{\left(r\left(1-\frac{A^*}{K}\right)-l\right)(A^*+a)}{g}$$

The expressions of the elements of the Jacobian matrix of the $RM_{(LV)}$ are (compare with S Eq. 13):

$$J_{(LV)1,1} = r_{lin} - g'Z'^* - l = r\left(1-\frac{A^*}{K}\right) - g\frac{Z^*}{(A^*+a)} - l = \frac{1}{A^*}\frac{dA^*}{dt} = 0$$

$$= J_{1,1} + r\frac{A^*}{K} + gZ^*\frac{a}{(A^*+a)^2} - g\frac{Z^*}{(A^*+a)}$$

$$J_{(LV)1,2} = -g_{lin}A^*_{(LV)} = -g\frac{A^*}{(A^*+a)} = J_{1,2} \quad \text{(S Eq. 21)}$$

$$J_{(LV)2,1} = eg_{lin}Z^*_{(LV)} = eg\frac{Z^*}{(A^*+a)} = J_{2,1}\frac{\boldsymbol{A^*+a}}{\boldsymbol{a}}$$

$$J_{(LV)2,2} = eg_{lin}A^*_{(LV)} - m = eg\frac{A^*}{(A^*+a)} - m = \frac{1}{Z^*}\frac{dZ^*}{dt} = 0 = J_{2,2}$$

**The Rosenzweig-MacArthur-Scheffer model - RMS**

The dynamics of the RMS model are given by:

$$\frac{dA}{dt} = rA\left(1-\frac{A}{K}\right) - gZ\frac{A}{A+a} - lA + lK \quad \text{(S Eq. 22)}$$

$$\frac{dZ}{dt} = egZ\frac{A}{A+a} - mZ - F\frac{Z^2}{Z^2+z^2}$$

The expressions of the elements of the Jacobian matrix of the RMS are:

$$J_{1,1} = r\left(1-\frac{A^*}{K}\right) - r\frac{A^*}{K} - gZ^*\frac{a}{(A+a)^2} - l \quad \text{(S Eq. 23)}$$

$$J_{1,2} = -g\left(\frac{A^*}{A^*+a}\right)$$

$$J_{2,1} = egZ^*\frac{a}{(A^*+a)^2}$$

$$J_{2,2} = eg\frac{A^*}{A^*+a} - m - 2FZ^*\frac{z^2}{(Z^{*2}+z^2)^2}$$

**The Rosenzweig-MacArthur-Scheffer model with RM dynamics – $RMS_{(RM)}$**

The dynamics of the $RMS_{(RM)}$ model are given by:

$$\frac{dA_{(RM)}}{dt} = rA_{(RM)}\left(1-\frac{A_{(RM)}}{K}\right) - gZ_{(RM)}\frac{A_{(RM)}}{A_{(RM)}+a} - lA_{(RM)} + lK$$

$$\frac{dZ_{(RM)}}{dt} = egZ_{(RM)}\frac{A_{(RM)}}{A_{(RM)}+a} - mZ_{(RM)} - F_{lin}Z_{(RM)} \quad \text{(S Eq. 24)}$$

with

$$F_{lin} = \frac{FZ^*}{(Z^{*2}+z^2)} \quad \text{(S Eq. 25)}$$

The expressions of the equilibria of the $RMS_{(RM)}$ model are:

$$A^*_{(RM)} = A^* \quad \text{(S Eq. 26)}$$

$$Z^*_{(RM)} = Z^*$$

The expressions of the elements of the Jacobian matrix of the $RMS_{(RM)}$ are (compare with S Eq. 23):

$$J_{(RM)1,1} = r\left(1 - \frac{A^*_{(RM)}}{K}\right) - r\frac{A^*_{(RM)}}{K} - gZ^*_{(RM)}\frac{a}{\left(A^*_{(RM)}+a\right)^2} - l = r\left(1-\frac{A^*}{K}\right) - r\frac{A^*}{K} - gZ^*\frac{a}{(A^*+a)^2} - l = J_{1,1}$$

$$J_{(RM)1,2} = -g\left(\frac{A^*_{(RM)}}{A^*_{(RM)}+a}\right) = -g\frac{A^*}{(A^*+a)} = J_{1,2} \quad \text{(S Eq. 27)}$$

$$J_{(RM)2,1} = egZ^*_{(RM)}\frac{a}{\left(A^*_{(RM)}+a\right)^2} = egZ^*\frac{a}{(A^*+a)^2} = J_{2,1}$$

$$J_{(RM)2,2} = eg\left(\frac{A^*_{(RM)}}{A^*_{(RM)}+a}\right) - m - F_{lin} = eg\frac{A^*}{(A^*+a)} - m - \frac{FZ^*}{(Z^{*2}+z^2)} = \frac{1}{Z^*}\frac{dZ^*}{dt} = 0$$
$$= J_{2,2} + 2FZ^*\frac{z^2}{(Z^{*2}+z^2)^2} - \frac{FZ^*}{(Z^{*2}+z^2)}$$

**The Rosenzweig-MacArthur-Scheffer model with Lotka-Volterra-Verhulst dynamics – $RM_{(LVV)}$**

The dynamics of the $RMS_{(LVV)}$ model are given by:

$$\frac{dA_{(LVV)}}{dt} = rA_{(LVV)}\left(1 - \frac{A_{(LVV)}}{K}\right) - g_{lin}Z_{(LVV)}A_{(LVV)} - lA_{(LVV)} + lK$$

$$\frac{dZ_{(LVV)}}{dt} = eg_{lin}Z_{(LVV)}A_{(LVV)} - mZ_{(LVV)} - F_{lin}Z_{(LVV)} \quad \text{(S Eq. 28)}$$

with

$$g_{lin} = \frac{g}{(A^*+a)} \quad \text{(S Eq. 29)}$$

$$F_{lin} = \frac{FZ^*}{(Z^{*2}+z^2)}$$

The expressions of the equilibria of the $RMS_{(LVV)}$ model are:

$$A^*_{(LVV)} = A^* \quad \text{(S Eq. 30)}$$

$$Z^*_{(LVV)} = Z^*$$

The expressions of the elements of the Jacobian matrix of the $RMS_{(LVV)}$ are (compare with S Eq. 23):

$$J_{(LVV)1,1} = r\left(1 - \frac{A^*_{(LVV)}}{K}\right) - r\frac{A^*_{(LVV)}}{K} - g_{lin}Z^*_{(LVV)} - l = r\left(1-\frac{A^*}{K}\right) - r\frac{A^*}{K} - g\frac{Z^*}{(A^*+a)} - l = J_{1,1} + gZ^*\frac{a}{(A^*+a)^2} - g\frac{Z^*}{(A^*+a)}$$

$$J_{(LVV)1,2} = -g_{lin}A^*_{(LVV)} = -g\frac{A^*}{(A^*+a)} = J_{1,2} \quad \text{(S Eq. 31)}$$

$$J_{(LVV)2,1} = eg_{lin}Z^*_{(LVV)} = eg\frac{Z^*}{(A^*+a)} = J_{2,1}\frac{A^*+a}{a}$$

$$J_{(LVV)2,2} = eg_{lin}A^*_{(LVV)} - m - F_{lin} = eg\frac{A^*}{(A^* + a)} - m - \frac{FZ^*}{(Z^{*2} + z^2)} = \frac{1}{Z^*}\frac{dZ^*}{dt} = 0$$

$$= J_{2,2} + 2FZ^*\frac{z^2}{(Z^{*2} + z^2)^2} - \frac{FZ^*}{(Z^{*2} + z^2)}$$

**The Rosenzweig-MacArthur-Scheffer model with LV dynamics - RMS(LV)**

The dynamics of the RMS(LV) model are given by:

$$\frac{dA_{(LV)}}{dt} = r_{lin}A_{(LV)} - g_{lin}Z_{(LV)}A_{(LV)} - lA_{(LV)} + lK$$

(S Eq. 32)

$$\frac{dZ_{(LV)}}{dt} = eg_{lin}Z_{(LV)}A_{(LV)} - mZ_{(LV)} - F_{lin}Z_{(LV)}$$

with

$$r_{lin} = r\left(1 - \frac{A^*}{K}\right) \qquad \text{(S Eq. 33)}$$

$$g_{lin} = \frac{g}{(A^*+a)}$$

$$F_{lin} = \frac{FZ^*}{(Z^{*2}+z^2)}$$

The expressions of the equilibria of the RMS(LV) model are:

$$A^*_{(LV)} = A^* \qquad \text{(S Eq. 34)}$$

$$Z^*_{(LV)} = Z^*$$

The expressions of the elements of the Jacobian matrix of the RMS(LV) are (compare with S Eq. 23):

$$J_{(LV)1,1} = r_{lin} - g_{lin}Z^*_{(LV)} - l = r\left(1 - \frac{A^*}{K}\right) - g\frac{Z^*}{(A^*+a)} - l = J_{1,1} + \boldsymbol{r}\frac{\boldsymbol{A^*}}{\boldsymbol{K}} + \boldsymbol{gZ^*}\frac{\boldsymbol{a}}{(\boldsymbol{A^*+a})^2} - \boldsymbol{g}\frac{\boldsymbol{Z^*}}{(\boldsymbol{A^*+a})}$$

$$J_{(LV)1,2} = -g_{lin}A^*_{(LV)} = -g\frac{A^*}{(A^*+a)} = J_{1,2} \qquad \text{(S Eq. 35)}$$

$$J_{(LV)2,1} = eg_{lin}Z^*_{(LV)} = eg\frac{Z^*}{(A^*+a)} = J_{2,1}\frac{\boldsymbol{A^*+a}}{\boldsymbol{a}}$$

$$J_{(LV)2,2} = eg_{lin}A^*_{(LV)} - m - F' = eg\frac{A^*}{(A^* + a)} - m - \frac{FZ^*}{(Z^{*2} + z^2)} = \frac{1}{Z^*}\frac{dZ^*}{dt} = 0$$

$$= J_{2,2} + \boldsymbol{2FZ^*}\frac{\boldsymbol{z^2}}{(\boldsymbol{Z^{*2}} + \boldsymbol{z^2})^2} - \frac{\boldsymbol{FZ^*}}{(\boldsymbol{Z^{*2}} + \boldsymbol{z^2})}$$
