## Supplemental Figure 1 for "Bridging theories for ecosystem stability through structural sensitivity analysis of ecological models in equilibrium"

**S2 Appendix**

**Results of the Rosenzweig-MacArthur-Scheffer model for increasing carrying capacity (*K*)**

The pattern that the increasing productivity of the system ends up only in increasing biomass of the consumer has been lost (S1 Fig 1a, line A*) and the model shows a more complex response of resource and consumer abundance to enrichment. In line with the LVV and RM model, the RMS model first goes through a transcritical bifurcation when increasing $K$ (S1 Fig 1a, $K = 1.0$), and shows a stable equilibrium thereafter. At a value of $K = 3.39$, however, a saddle-node bifurcation occurs and the equilibrium densities of resources and consumers suddenly switch to a different level, lower for the resource and higher for the zooplankton (S1 Fig 1a). At decreasing carrying capacity, this switch takes place at a lower value of $K = 3.12$ and hence the system shows hysteresis and alternative stable states also for K. In addition to these alternative stable states, the system shows a supercritical Hopf bifurcation at $K = 3.89$ and the model shows stable limit cycles as its dynamic behavior at higher carrying capacities (S1 Fig 1b). For the linearized RMS model version $RMS_{(LV)}$, when increasing $K$, the model does show the transcritical bifurcation at the value of 1.0 but thereafter only shows stable focus for higher values of the carrying capacity (S1 Fig 1b).

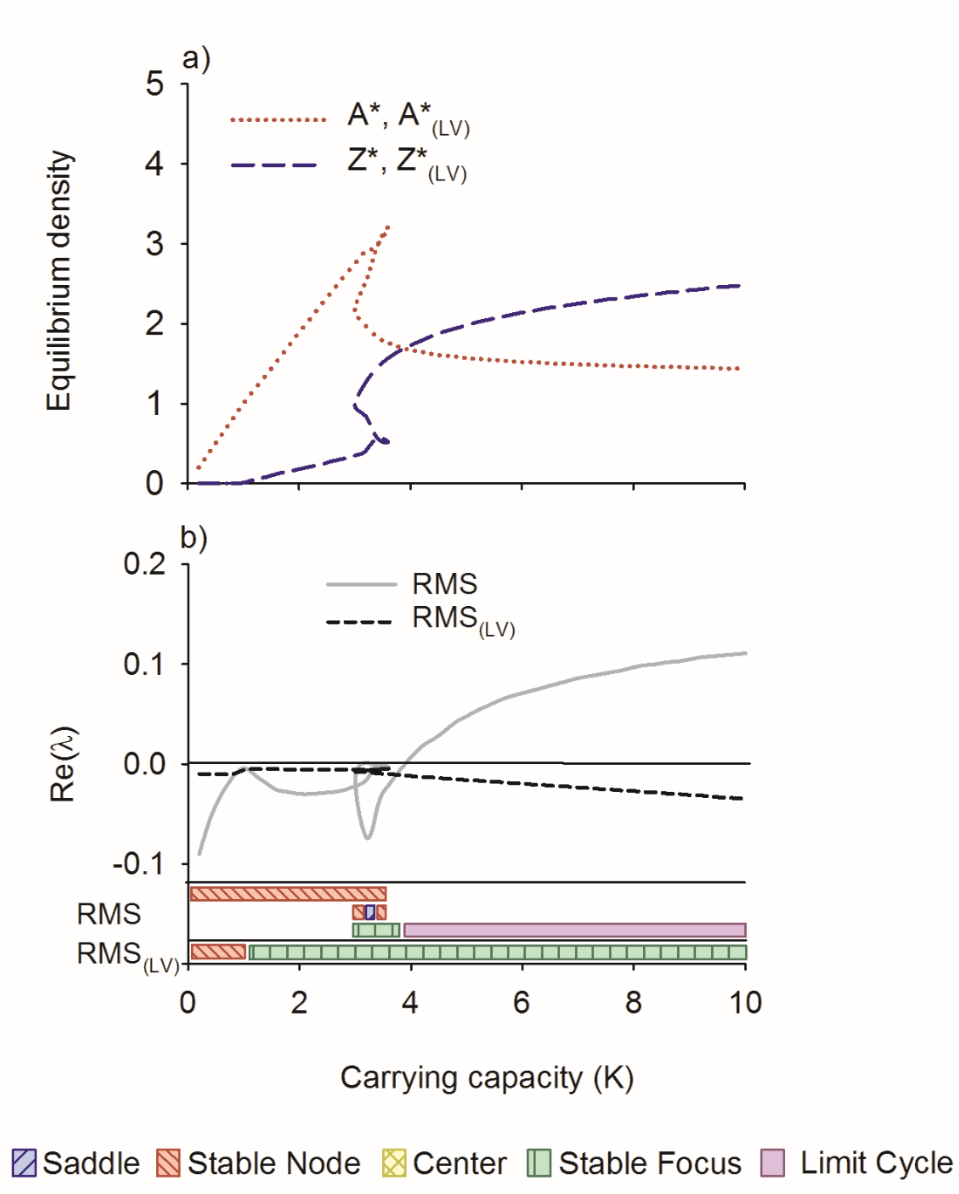


**S1 Fig 1.** Equilibrium densities (a) and the maximum real part of the eigenvalues (b) of the original Rosenzweig-MacArthur-Scheffer (RMS) and the linearized ($RMS_{(LV)}$) model along a gradient of increasing carrying capacity (*K*). Also the mode of behavior along the gradient is depicted. Parameters are $r = 0.5$, $g = 0.4$, $a = 0.6$, $l = 0.01$, $e = 0.6$, $m = 0.15$, $F = 0.05$ and $z = 0.5$.
